# Expanding gene families helps generate the metabolic robustness required for antibiotic biosynthesis

**DOI:** 10.1101/119354

**Authors:** Jana K Schniete, Pablo Cruz-Morales, Nelly Selem, Lorena T. Fernández-Martínez, Iain S Hunter, Francisco Barona-Gómez, Paul A Hoskisson

**Affiliations:** Strathclyde Institute of Pharmacy and Biomedical Sciences, University of Strathclyde, 161 Cathedral Street, Glasgow, G4 0RE, United Kingdom; Evolution of Metabolic Diversity Laboratory, Langebio, Cinvestav-IPN, Libramiento Norte Carretera Leon Km 9.6, 36821, Irapuato, Guanajuato, México; Department of Biology, Edgehill University, St Helens Road, Ormskirk, Lancashire, L39 4QP, UK

## Abstract

Expanding the genetic repertoire of an organism by gene duplication or horizontal gene transfer (HGT) can aid adaptation. *Streptomyces* species are prolific producers of bioactive specialised metabolites with adaptive functions in nature and some have found utility in human medicine such as antibiotics. Whilst the biosynthesis of these specialised metabolites is directed by dedicated biosynthetic gene clusters (BGCs), little attention has been focussed on how these organisms have evolved robustness into their genomes to facilitate the metabolic plasticity required to provide chemical precursors for biosynthesis. Here we show that specific expansions of gene families in central carbon metabolism have evolved and become fixed in *Streptomyces* bacteria to enable plasticity and robustness that maintain cell functionality whilst costly specialised metabolites are produced. These expanded gene families, in addition to being a metabolic adaptation, make excellent targets for metabolic engineering of industrial specialised metabolite producing bacteria.

## Introduction

A remarkable feature of specialised metabolite producing Actinobacterial genomes is the annotation of multiple genes that encode the same putative biochemical function^1,2^. This expansion of gene families by gene duplication or HGT is thought to introduce robustness into biological systems, which in turn facilitates evolvability and adaptation^3–5^. The expansion of gene families results in relaxed selection following the gene duplication or HGT event, that allows the accumulation of mutations which enable diversification of function to occur^6^. This suggests that gene family expansion within genomes is a key driver of biological innovation by facilitating adaptation^7^. The production of extensive specialised metabolites by certain Actinobacterial lineages is thought to be a key adaptive response to life in complex, highly competitive environments such as soil ^8–10^ and, as such, may drive the expansion of primary metabolic capability providing the metabolic robustness that facilitates the evolution of novel biosynthetic functions.

Surprisingly many central metabolic enzymes are non-essential for survival due to genetic redundancy through the presence of isoenzymes or alternative reactions. The redundancy allows cells to adapt to a variety of habitats and dynamic environmental conditions through provision of metabolic plasticity^11^. Whilst this has been studied in the unicellular enteric bacterium *Escherichia coli* and the yeast *Saccharomyces cerevisae*^12^, little attention has been paid to organisms with extensive specialised metabolism. Gene families of Actinobacterial developmental genes have been studied at the genetic level^7,13,14^ but little attention has been paid to either primary or specialised metabolism^15^ and how the supply of biosynthetic precursors is maintained during the adaptive response under challenging environmental conditions.

In Actinobacteria, production of specialised metabolites is frequently growth phase dependent and usually in response to nutrient starvation and during entry into sporulation^16^. This creates a potential metabolic conflict for an organism, where declining availability of metabolites may constrain certain cellular process in favour of others, such as reducing cellular pools of metabolites that are used directly for specialised metabolites. Under these conditions it is likely that genetic redundancy can promote robustness and plasticity that helps to maintain cellular function in the face of perturbation^17,18^.

Here we systematically examine the genetic redundancy within the genomes of specialised metabolite producing Actinobacteria to understand how genetic robustness enables the evolution of extensive specialized metabolism. Moreover, a detailed functional analysis of a redundant pyruvate kinase gene pair from *Streptomyces coelicolor* A3(2) indicates that biochemical diversification at the enzyme level facilitates the evolution of distinct physiological roles which enables functionality during the metabolic reprogramming that is associated with physiological differentiation.

## Results

### Gene expansion events are overrepresented in specialised metabolite producing Actinobacteria

To determine if gene expansion events in central metabolism occur with greater frequency in specialised metabolite producing organisms, a database of 614 Actinobacterial genomes spanning 80 genera was compiled. All genomes were retrieved from GenBank, and reannotated with RAST^19^ to ensure consistency of annotation across the database and were then analysed in a bespoke bioinformatics pipeline based on EvoMining^20^ (Supplementary Fig. S1). It was hypothesised that, if precursor supplying pathways are a contributing factor to the adaptive response of specialised metabolite production, then the enzymatic nodes contributing to precursor supply should be overrepresented in the database (Table 1 and Supplementary Table S1).

**Table 1.**
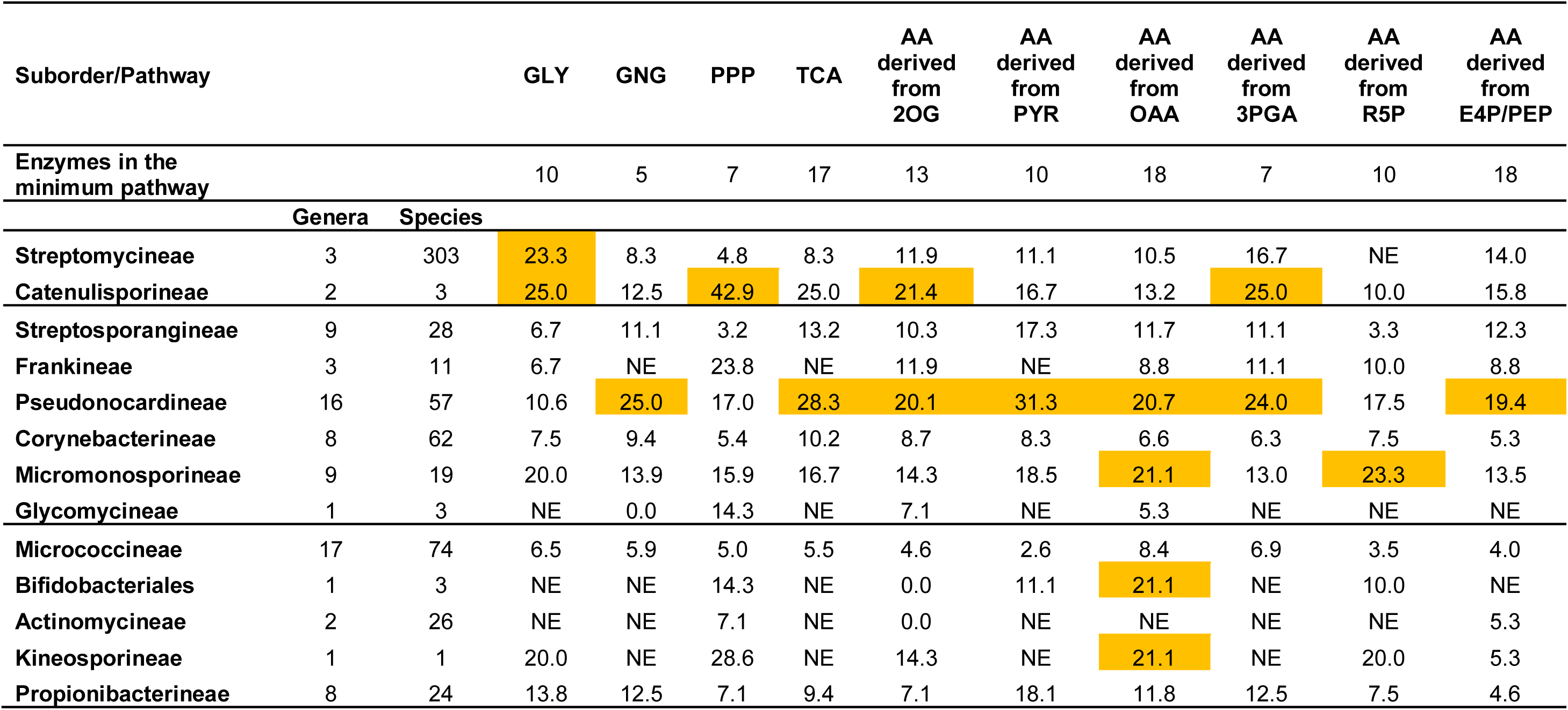
Percentage of primary metabolic pathway gene expansion per suborder and pathways of central carbon metabolism. (highest percentage of gene expansion for each pathway is highlighted in yellow). **Legend:** GLY = glycolysis, GNG = gluconeogenesis, PPP = pentose phosphate pathway, TCA = tricarboxylic acid cycle, AA = amino acids, 2OG = 2-oxo-glutarate, PYR = pyruvate, OAA = oxaloacetate, 3PGA = 3-phosphoglycerate, R5P = ribose-5-phosphate, E4P = erythrose-4-phosphate, PEP = phosphoenolpyruvate, NE = no expansion.

| Suborder/Pathway |  |  | GLY | GNG | PPP | TCA | AA<br>derived<br>from<br>2OG | AA<br>derived<br>from<br>PYR | AA<br>derived<br>from<br>OAA | AA<br>derived<br>from<br>3PGA | AA<br>derived<br>from<br>R5P | AA<br>derived<br>from<br>E4P/PEP |
| --- | --- | --- | --- | --- | --- | --- | --- | --- | --- | --- | --- | --- |
| Enzymes in the<br>minimum pathway |  |  | 10 | 5 | 7 | 17 | 13 | 10 | 18 | 7 | 10 | 18 |
|  | Genera | Species |  |  |  |  |  |  |  |  |  |  |
| Streptomycineae | 3 | 303 | 23.3 | 8.3 | 4.8 | 8.3 | 11.9 | 11.1 | 10.5 | 16.7 | NE | 14.0 |
| Catenulisporineae | 2 | 3 | 25.0 | 12.5 | 42.9 | 25.0 | 21.4 | 16.7 | 13.2 | 25.0 | 10.0 | 15.8 |
| Streptosporangineae | 9 | 28 | 6.7 | 11.1 | 3.2 | 13.2 | 10.3 | 17.3 | 11.7 | 11.1 | 3.3 | 12.3 |
| Frankineae | 3 | 11 | 6.7 | NE | 23.8 | NE | 11.9 | NE | 8.8 | 11.1 | 10.0 | 8.8 |
| Pseudonocardineae | 16 | 57 | 10.6 | 25.0 | 17.0 | 28.3 | 20.1 | 31.3 | 20.7 | 24.0 | 17.5 | 19.4 |
| Corynebacterineae | 8 | 62 | 7.5 | 9.4 | 5.4 | 10.2 | 8.7 | 8.3 | 6.6 | 6.3 | 7.5 | 5.3 |
| Micromonosporineae | 9 | 19 | 20.0 | 13.9 | 15.9 | 16.7 | 14.3 | 18.5 | 21.1 | 13.0 | 23.3 | 13.5 |
| Glycomycineae | 1 | 3 | NE | 0.0 | 14.3 | NE | 7.1 | NE | 5.3 | NE | NE | NE |
| Micrococcineae | 17 | 74 | 6.5 | 5.9 | 5.0 | 5.5 | 4.6 | 2.6 | 8.4 | 6.9 | 3.5 | 4.0 |
| Bifidobacteriales | 1 | 3 | NE | NE | 14.3 | NE | 0.0 | 11.1 | 21.1 | NE | 10.0 | NE |
| Actinomycineae | 2 | 26 | NE | NE | 7.1 | NE | 0.0 | NE | NE | NE | NE | 5.3 |
| Kineosporineae | 1 | 1 | 20.0 | NE | 28.6 | NE | 14.3 | NE | 21.1 | NE | 20.0 | 5.3 |
| Propionibacterineae | 8 | 24 | 13.8 | 12.5 | 7.1 | 9.4 | 7.1 | 18.1 | 11.8 | 12.5 | 7.5 | 4.6 |

Expansion events were defined as cases where the number of enzyme family members per suborder had a value equal or higher than the mean number of members per phylum plus its standard deviation. The glycolytic pathway showed highest number of gene expansion events, in the Streptomycineae and Catenulisporineae with 23.3% and 25.0% more genes encoding glycolytic function than was average for that pathway in the phylum Actinobacteria respectively (Table 1). Pseudonocardineae showed highest number of gene expansions in gluconeogenesis (25% higher than the mean phylum value) and in the TCA cycle (28.3% higher than the mean phylum value). This was also true for many amino acid biosynthetic pathways. Where the main precursor is derived from 2-oxo-glutarate (Glu, Gln, Pro, Arg), expansion was 20.1% more than the mean suborder value, with pyruvate derived amino acids (Ala, Ile, Leu, Val; 31.3%), oxaloacetate derived amino acids (Asp, Asn, Thr, Met, Lys; 20.7%), 3-PGA derived amino acids (Gly, Ser, Cys; 24%) and E4P/PEP derived amino acids (Tyr, Phe, Trp; 19.4%).

Focusing on the genus *Streptomyces,* which is renowned as being amongst the most talented of genera in terms of specialised metabolite production, it was found that 14 enzymatic steps from central metabolism (Glycolysis, TCA cycle, and amino acid metabolism) represented gene expansion events, such that they are overrepresented in this genus compared to the rest of the database. The following enzyme functions were found to be over-represented in the genus *Streptomyces* compared to the whole Actinobacterial phylum: phosphofructokinase (PFK), pyruvate kinase (PK), pyruvate phosphate dikinase (PPDK), malic enzyme (ME), pyruvate dehydrogenase complex E1 (PDHC E1), chorismate mutase, acetylglutamate kinase, diaminopimelate decarboxylase, aspartate aminotransferase, aspartate-semialdehyde dehydrogenase, serine hydroxymethyltransferase, glutamine synthetase, arginiosuccinate lyase and methionine synthetase (Table S1). To investigate how gene expansions in Actinobacteria are a potential prerequisite for increasing robustness in specialised metabolism capability, the two pyruvate kinases kinases from *Streptomyces* were studied further due to their central role in carbon metabolism linking glycolysis, gluconeogenesis and the TCA cycle.

### Pyruvate kinases in *Streptomyces* arose by gene duplication

PK catalyses the terminal step of glycolysis, converting one molecule of phosphoenolpyruvate to one molecule of pyruvate using ADP as the phosphor-acceptor resulting in the production of ATP. PK therefore plays a key role in linking glycolysis and the citric acid cycle. Moreover, it results in the formation of a direct precursor of Acetyl-CoA, which feeds directly into polyketide specialised metabolites. A high resolution species level phylogeny of the Actinobacteria was constructed using the β-subunit of RNA polymerase (RpoB)^21,22^ which allowed the segregation of the phylum into three distinct phylogenetic branches: one composed of Streptomycineae and Catenulisporinae, a second including Propionibacterineae, Actinomycineae, Bifidobacteriales and Micrococcineae and the third one with Micromonosporineae, Glycomycineae, Corynebacterineae, Pseudonocardineae, Frankineae and Streptosporangineae (Fig. 1A).

**Figure 1:**
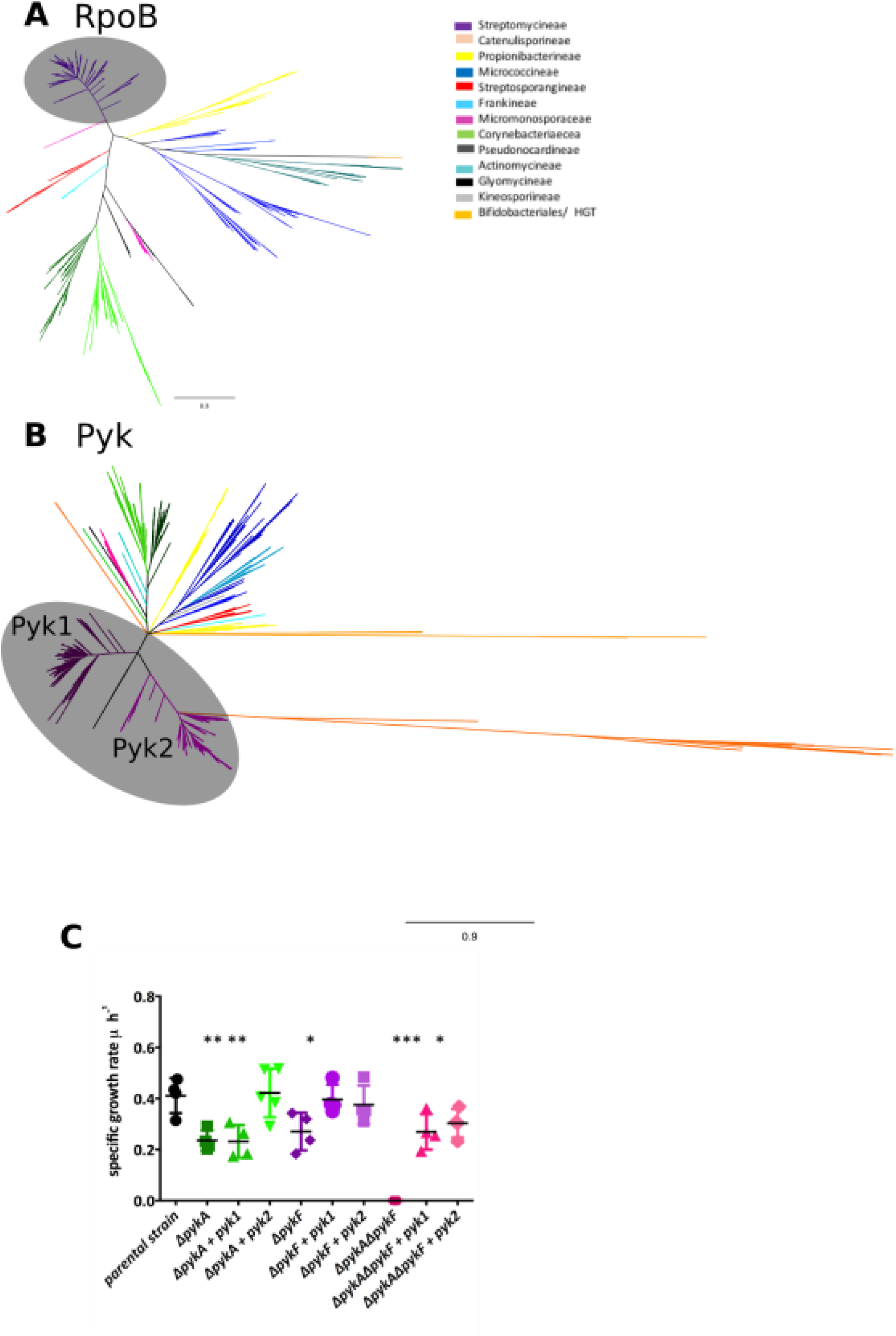
Phylogenetic Analysis of RpoB and PK across 80 different actinobacterial genera grouped and colour coded by family **(A)** Phylogenetic tree based on RpoB protein sequences **(B)** Phylogenetic tree based on pyruvate kinase protein sequences. Grey circles indicate the *Streptomycineae* **(C)** Specific growth rate of *E. coli* pyruvate kinase mutants (Δ*pykA*, Δ*pykF*, Δ*pykA*Δ*pykF*) and the complemented mutants with either *pyk1* or *pyk2* from *S. coelicolor* on M9 Medium with glucose as carbon source. * *p*-value < 0.05 ** *p*-value < 0.01 *** *p*-value ≤ 0.001

A second phylogeny of the annotated PKs of the Actinobacteria was constructed. It indicated that there is a high level of congruence with the RpoB phylogeny as expected for a central metabolic enzyme (Fig. 1B). However, a bifurcating topology within the Streptomycineae family was observed, which contained the two genes encoding the putative PKs. This topology indicates that a gene duplication event occurred, which gave rise to two PKs within this group. Analysis of 286 *Streptomyces* species showed that 281 species have duplicate copies of PK, three species possess a single copy (*S. somaliensis, S.* sp NRRL F5135 and *S. scrabrisporus),* two species have three copies (*S. olindensis* and *S. sp.* AcH505) and a single species has four copies (*S. resistomycificus).* Interestingly, *S. sp* AcH505 and *S. resistomycificus* had one copy of *pyk* in each main branch of the PK tree and additional copies were found to be phylogenetically distant, suggesting that these copies were acquired through horizontal gene transfer (HGT). Overall, 92 % (302 of 327) Actinobacterial genomes outside of the genus *Streptomyces* encoded a single PK reinforcing the uniqueness of the duplication in this genus (Fig. 1B).

To determine if the duplicate PKs annotated in the *Streptomyces* genome have pyruvate kinase activity we used the two PKs from the model streptomycete, *S. coelicolor* A3(2), in genetic complementation tests of PK mutants of *Escherichia coli. E. coli* also has two PKs: a primary enzyme *pykF,* which is a Type I enzyme, regulated allosterically by fructose 1,6 biphosphate (FBP) and a distinct secondary Type II PK (*pykA*), regulated allosterically by AMP^23^. In *Streptomyces,* both PKs (Pyk1 and Pyk2) are Type I enzymes, homologous to PykF of *E. coli* (40.6 % and 41.3 % identity respectively). To test for functional complementation, *E. coli* mutants (Δ*pykA*, Δ*pykF* and a Δ*pykA* Δ*pykF* double mutant, Table S3) were tested, along with the isogenic parental strain (*E. coli* BW25113) for their ability to grow under a range of physiological conditions (Fig. 1C). In LB (for which PK is dispensable for growth) and M9 plus acetate as the sole carbon source (where PK is also dispensable for growth), little difference was observed in the specific growth rate (h^-1^) of the strains (Data not shown). When the strains were grown in M9 plus glucose as the sole carbon source (where PK is essential for growth) the *E. coli ΔpykA ΔpykF* double mutant was unable to grow, but could be genetically complemented with either *pyk1* (SCO2014) or *pyk2* (SCO5423) from *S. coelicolor.* The individual *E. coli ΔpykA* and the *ΔpykF* mutants had reduced specific growth rates (around 50% of the isogenic parent strain) in M9 plus glucose. Genetic complementation with *pyk1* or *pyk2* from *S. coelicolor* was able to fully restore growth of an *E. coli ΔpykF* mutant as expected. The *E. coli ΔpykA* mutant could only be complemented with *pyk1* from *S. coelicolor,* suggesting a much more limited physiological role for *pykA* in *E. coli* (Fig. 1C). These data confirm that both *pyk1* and *pyk2* from *S. coelicolor* have retained PK activity following the duplication event but suggests that each has diverged and evolved different physiological roles.

Given that the PKs in *S. coelicolor* have diverged following duplication, we assessed the level of selection imposed on the PKs of *Steptomyces* by calculating the ratio of non-synonymous changes (dN) to synonymous changes (dS). Twenty PK sequences from 10 *Streptomyces* genomes were chosen to calculate the dN, dS and dN/dS values. The dN/dS ratio for pairs of *pyk* sequences for each of the genomes yielded dN/dS ratios ranging from 0.407 to 0.500, suggesting that PKs in *Streptomyces* are under strong purifying selection (Table. S2). Such high levels of purifying selection indicate that the duplication event in *Streptomyces* is likely to be ancient and is consistent with the PK tree topology (Fig. 1B).

### The two pyruvate kinases in *Streptomyces* have distinct physiological roles

To determine the roles played by the PKs in growth, development and antibiotic production, a series of mutant *S. coelicolor* strains was constructed and genetically complemented (Table S3, S4 & Fig. S2). Deletion mutants and transposon insertion mutants showed similar phenotypes (Fig.S2) and all subsequent work was carried out with transposon insertion mutants. Growth on nutrient agar showed no differences between the strains, except when an additional copy of *pyk1* was present in WT *in trans* (Fig. 2A). During culture on solid minimal medium with 1% glucose as carbon source, the strains showed no growth defects when compared to wild-type (Fig. 2A). Interestingly, the *pyk1::Tn5062* mutant showed an increase in specialised metabolite production (Fig. 2A, 2C & 2D). The strain *pyk2*::Tn5062 was marginally affected in growth and showed no over expression of specialised metabolites (Fig. 2A). No changes in growth rate were observed in rich medium (YEME medium) for the WT, mutants or complemented strains (Fig. 2B). However, growth of the strains in this medium showed an increase in production of coelimycin^24^ and undecylprodigiosin (RED) in the *pyk1::Tn5062* mutant (Fig. 2C) whereas a *pyk2::Tn5062* mutant showed reduced antibiotic yields (Fig. 2C & 2D). These data suggest that each PK isoenzyme plays a distinct physiological role in growth of *Streptomyces* and perturbation of central metabolism by their deletion or addition affects specialised metabolite biosynthesis. This would suggest that either the PKs are transcriptionally regulated to be expressed at key stages of the *Streptomyces* lifecycle or are regulated at the post-transcriptional/translational level. To test this hypothesis, we used semi-quantitative RT-PCR to examine the expression of the PKs from *S. coelicolor* throughout growth, relative to the multiplexed vegetative sigma factor *hrdB.* We found that both genes were constitutively expressed throughout growth (vegetative hyphae, aerial hyphae and during sporulation) relative to *hrdB* (Fig. 3A). To further characterise transcription, we used quantitative qRT-PCR at two time points during log and stationary phase during either glycolytic growth (glucose as sole carbon source) or gluconeogenic growth (Tween as sole carbon source) to analyse the expression levels of *pyk1*, *pyk2* and *hrdB.* Normalising expression to *hrdB,* there was an expected decrease in *pyk1* expression on Tween compared to glucose during the log and stationary phases of growth (3-fold and 8-fold respectively; Fig. 3A and B). Comparison of *pyk1* and *pyk2* during the logarithmic growth phase indicated that *pyk2* had a 1.5-fold lower level of expression than in stationary phase versus log phase when grown on Tween as the sole carbon source, with all other conditions showing no significant changes in expression between *pyk1* and *pyk2* (Fig. 3Bi), suggesting that activity of PKs in *Streptomyces* is likely to be controlled at the post-translational level. Expression of *pyk2* also showed a decrease in expression on Tween compared to glucose during both phases, but the change was not significant (Fig 3Bii).

**Figure 2:**
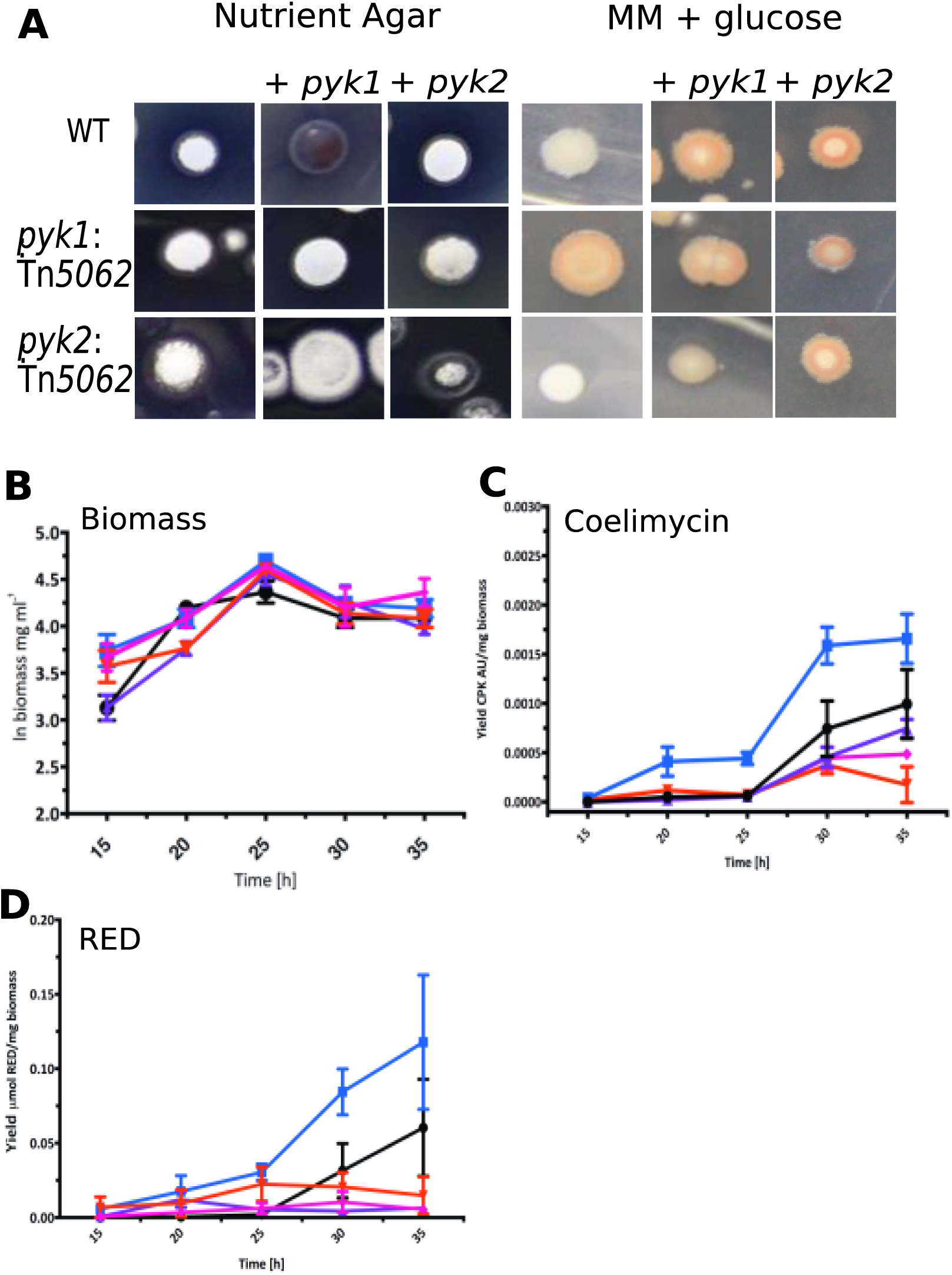
Phenotypic characterisation of pyruvate kinase mutants **(A)** Wild-Type and *pkyl* & *pyk2* transposon mutants of *S. coelicolor* grown on nutrient agar and minimal medium with glucose, with complemented strains with either *pyk1* or *pyk2 in trans* and WT strains with additional copies of *pyk1* or *pyk2* **(B)** Growth curve of *S. coelicolor* WT, pyruvate kinase mutants and complemented strains in liquid YEME medium **(C)** coelimycin production yield (absorption unit/mg biomass) and **(D)** undecylprodigiosin (RED) yield during growth in YEME medium. WT (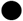) and pyruvate kinase mutants (*pyk1* 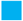; *pyk2* 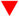) and complemented strains (*pyk1* complemented 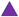; *pyk2* complemented 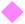).

**Figure 3:**
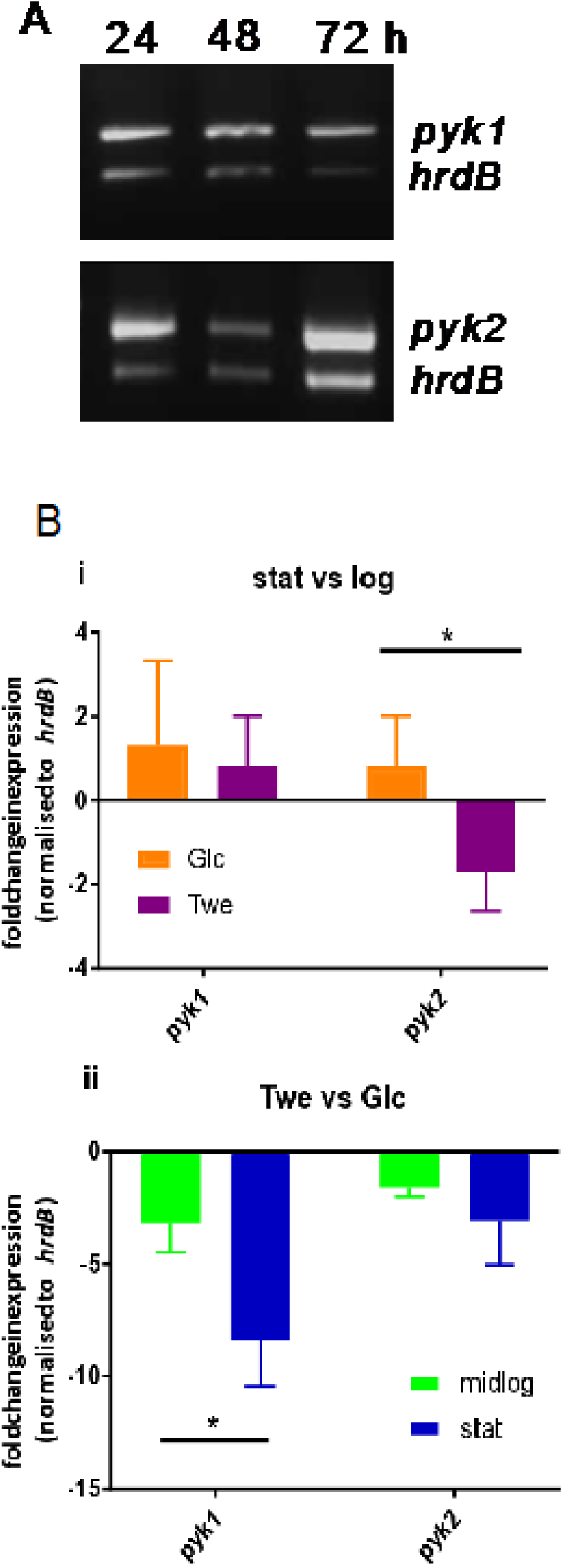
(**A**) Semi-quantitative RT-PCR of expression of *pyk1, pyk2 and hrdB* throughout the lifecycle of *Streptomyces coelicolor.* **(B)** Fold change expression of *pyk1* and *pyk2* normalised to *hrdB* expression from three biological replicates measuring expression levels by qPCR on growth in minimal medium with either glucose or tween as carbon source during log or stationary phase comparing expression (i) stationary phase versus log phase and (ii) Tween versus glucose. * p-value < 0.05

### Pyk1 and Pyk2 in *Streptomyces* have key substrate affinity differences and specific effector molecules

In order to understand the biochemical control of the two PKs of *S. coelicolor,* we purified each enzyme and studied their biochemical characteristics. Both Pyk1 and Pyk2 were activated by the effector molecule AMP and they both showed Michaelis-Menten type kinetics for the substrate ADP. For Pyk1, S_0.5_ was 4-fold lower in the presence of 1 mM AMP (0.59 mM down to 0.15 mM; Table 2), while V_max_ also increased 3.5-fold (from 21 U/mg to 73.3 U/mg). Pyk2 showed a five-fold increase of V_max_ in the presence of 1 mM AMP (from 1.2 U/mg to 6.7 U/mg).

**Table 2:** Kinetic characteristics of Pyk1 and Pyk2 for the substrates ADP, PEP and the activator AMP

| Substrate | Parameter | no AMP |  | 1 mM AMP |  |
| --- | --- | --- | --- | --- | --- |
|  |  | Pyk1 | Pyk2 | Pyk1 | Pyk2 |
| 5 mM PEP<br>ADP | $V_{\max}$ (U/mg) | 21.0 | 1.2 | 73.3 | 6.7 |
| | $S_{0.5}$ (mM) | 0.6 | 0.3 | 0.2 | 0.1 |
| | $k_{\text{cat}}$ (sec<sup>-1</sup>) | 941 | 39 | 4703 | 215 |
| | $k_{\text{cat}}/K_m$ (mM<sup>-1</sup> s<sup>-1</sup>) | 1594 | 145 | 31359 | 2388 |
| 1.5 mM ADP<br>PEP | $V_{\max}$ (U/mg) | 14.1 | 0.5 | 65.5 | 9.1 |
| | $S_{0.5}$ (mM) | 3.5 | 1.3 | 1.1 | 8.6 |
|  | Hill Coefficient | 3.7 | 1.5 | 1.8 | 7.1 |
| | $k_{\text{cat}}$ (sec<sup>-1</sup>) | 350 | 18 | 4200 | 336 |
| | $k_{\text{cat}}/S_{0.5}$ (mM<sup>-1</sup> s<sup>-1</sup>) | 100 | 14 | 4000 | 39 |
| AMP | $V_{\max}$ (U/mg) | 8.2 | 1 | | |
| | $S_{0.5}$ (mM) | 0.01 | 3.8 | | |
| | $k_{\text{cat}}$ (sec<sup>-1</sup>) | 423.7 | 42.0 | | |
| | $K_{\text{cat}}/S_{0.5}$ (mM<sup>-1</sup> s<sup>-1</sup>) | 42373 | 11 | | |

S_0.5_ decreased three-fold (from 0.27 mM to 0.09 mM; Table 2, Fig. S2). There were profound differences in the PEP kinetics for both PKs, with both isoenzymes demonstrating Hill-type cooperative binding kinetics with AMP. In the presence of 1 mM AMP, V_max_ of Pyk1 increased five-fold (14.05 U/mg to 65.45 U/mg), S0.5 decreased more than three-fold (3.49 to 1.05 mM) and the Hill coefficient was approximately halved (from 3.7 to 1.8). For Pyk2 in the presence of 1 mM AMP, V_max_ was 9.1 U/mg compared to 0.5 U/mg without AMP, with S0.5 increased from 1.3 mM to 8.6 mM. Under these conditions the Hill coefficient increased from 1.45 to 7.1 (Table 2). Further analysis demonstrated that Pyk1 has a much higher affinity for AMP (S_0.5_ = 0.01 mM), compared to Pyk2 (S0.5 = 3.8 mM), with a concomitant increase in V_max_ (8.2 U/mg for Pyk1 compared to 1 U/mg for Pyk2, Fig S3C). The turnover rate constant (K_cat_) for Pyk1 was >20 fold greater (4703 sec^-1^) than that of Pyk2 (215 sec^-1^; Table 2). Interestingly Pyk1 was also shown to be highly stimulated by ribose-5-phosphate (Fig. S3). Intriguingly it is known that flux through the pentose phosphate pathway increases during entry into stationary phase in streptomycetes ^25^ suggesting that Pyk1 activity is stimulated during periods of starvation and during antibiotic production to rebalance reduced glycolytic flux and entry of substrates in to the TCA cycle.

## Discussion

Metabolic robustness is a key biological adaptation to coping with environmental perturbation that enables an organism to persist in a given environment^17^. The ability of an organism to adapt to dynamic and competitive environments requires minimization of the effects of metabolic perturbations, which can be achieved through gene family expansion either following gene duplication or HGT. During nutrient stress, intracellular concentrations of key metabolic intermediates can fall which, when coupled with increasing demand during specialized metabolite biosynthesis, may produce metabolic conflict in cells. PK occupies a key position in carbon metabolism bridging glycolysis to the TCA cycle and plays a central role in the generation of ATP and precursors for the synthesis of specialised metabolite precursors (Acetyl-CoA, amino acids, organic acids etc)^26^. Duplication may promote adaptation and metabolic robustness through the evolution of altered substrate affinity and enzyme efficiency, which in turn enables key cellular process to proceed during times of perturbation, enabling specialised metabolite production and cellular growth to proceed, as we have demonstrated here.

Enzyme family expansion in central metabolism is widespread in the Actinobacteria with the Streptomycineae showing extensive enzyme family expansion in glycolysis. The duplication of pyruvate kinase is ancient, and has permitted subsequent divergence of the gene pair to evolve distinct physiological roles, where Pyk2 appears to function as a house-keeping PK, with a higher affinity for PEP (when AMP is low) and Pyk1 exhibits strong activation as AMP concentrations rise. An increased AMP concentration is a well-established starvation signal in bacteria^27^ and may serve to increase flux through the terminal end of glycolysis during starvation to facilitate precursor supply for specialised metabolites. Moreover the PPP intermediate, ribose-5-phosphate, stimulates Pyk1 activity providing a physiological link between the PPP and the associated generation of NADPH which has established links to specialised metabolite synthesis, including those overproduced by strains engineered in this work^25^. Disruption of *pyk1* in *S. coelicolor* lead to increased levels of coelimycin and undecylprodigiosin when grown in rich medium (such as in industrial situations), but no significant increase in the yield of the polyketide antibiotic actinorhodin, which may suggest different metabolic control points affect biosynthesis of chemically similar specialised metabolites. We also demonstrate that the duplication of the primary metabolic enzyme, PK promotes metabolic robustness and influences the production of specialised metabolites. Understanding the evolution of central metabolism in conjunction with specialised metabolism can contribute to our fundamental understanding of the ability of Actinobacteria to produce a plethora of useful molecules and can help inform on novel approaches to metabolic engineering.

## Methods

### Database generation and bioinformatics analysis

The NCBI database (http://www.ncbi.nlm.nih.gov/genbank/wgs) was the source of actinobacterial genomes having a minimum coverage of 25x and less than 30 contigs per Mbp. To ensure a wide range of phylogeny, a selection of 614 species from 80 genera were included. Each genome was re-annotated using RAST^19^ and the annotation files used to determine the frequency of each functional annotation. The mean of occurrences of each functional role was calculated per genus and examples which had a value equal or higher than the mean plus its standard deviation were defined as a ‘gene expansion event’. Each candidate protein sequence was extracted from the actinobacterial genome database to form a BLASTP analysis working database. The sequences were then aligned with MUSCLE V3.8.31^28^, alignments were scrutinised using Jalview V2.10.1^30^ to ensure that at least 25 % coverage with the query was achieved; if not sequences and expansions were discarded from the working database. Phylogenetic analysis of the alignments was conducted using MrBayes^29^ V3.2.3 with trees visualised in FigTree V1.4.2(http://tree.bio.ed.ac.uk/software/figtree/). Sequences obtained from NCBI gene database and aligned with ClustalW algorithm in MEGA V6.06 ^30^ and the synonymous (dS) and nonsynonymous (dN) changes were determined using the Distance Model Function with Nei-Gojobori^31^ with Jukes-Cantor algorithm^32^ Bootstrap.

### Growth and mutant construction in *Streptomyces*

Routine growth, spore generation and conjugation of *Streptomyces* strains were carried out according to Kieser *et al.,*^33^ Antibiotic titres were determined according to Gottelt et al ^24^ for coelimycin and Kieser et al.,^33^ for RED and ACT. *S. coelicolor* gene knock-out mutants (*Δpyk1* and *Δpyk2)* were constructed using PCR-targeted gene replacement with an apramycin resistance cassette *(acc(3)IV)* using the Redirect system^34^ and the primers reported in Table S5. *S. coelicolor* transposon insertion mutants *(pyk1::Tn5062* and pyk2::Tn5062) were constructed using Tn5062 mutagenised cosmids as described in Fernández-Martínez et al ^35^. Each cosmid was first verified by restriction analysis before being conjugated into *Streptomyces.* All strains were verified by PCR and sequencing of the respective products.

### Interspecies complementation

*E. coli* single mutants were from the Keio collection^36^ of *E. coli* BW25113^36^ and the double mutant was constructed using Lambda Red recombination of *pykF* according to Datsenko and Wanner^37^. Complementation studies used the *Streptomyces pyk1* and *pyk2* cloned into pET100_TOPO (Invitrogen).

*E. coli* growth curves were carried out in 250 ml flasks with a working volume of 50 ml of either LB or M9 medium with 1% (w/v) glucose or 0.4% sodium acetate (w/v) as carbon source. Flasks were inoculated from an overnight culture (1% v/v) including the appropriate antibiotics and 1 mM IPTG to induce expression of the PKs. Growth was followed at OD600 at 37°C with shaking at 250 rpm. The specific growth rate was determined from the semi-logarithmic plot of biomass concentration.

### Protein overexpression and purification

The coding sequence of *pyk1* was codon optimised for *E coli* and amplified from the vector pEX-K4 using the primers in Table 2. The native version of *pyk2* was used to amplify the coding sequence using the primers in Table 2. Both coding sequences were cloned into the pET100 TOPO vector (Invitrogen) according to the manufacturer’s instructions. Overexpression of *pyk1* was in *E. coli* Origami B on LB with 1 % (w/v) glucose at 30 °C until an OD_600_ of 0.4 was reached and the expression was induced with 0.05 mM IPTG at 18°C overnight. Pyk2 overexpression was in *E. coli* Rosetta using Auto Induction medium (component per L: 10 g tryptone, 5 g yeast extract, 3.3 g (NH_4_)_2_SO_4_, 6.8 g KH_2_PO_4_, 7.1 g Na_2_HPO_4_, 0.5 g Glucose, 2.0 g α-Lactose, 0.15 g MgSO_4_) grown at 37°C for 2 h and then reduced to 18°C for overnight cultivation. Cells were disrupted by sonication. Pyk1 and Pyk2 were purified by nickel affinity chromatography using HisTrap TM FF crude (GE Healthcare) with Binding Buffer (100 mM KH_2_PO_4_ pH7.2, 10% glycerol (v/v), 100 mM NaCl, 20 mM imidazole). Tagged proteins were eluted with increasing imidazole concentration (Elution buffer: 100 mM KH_2_PO_4_ pH7.2, 10% glycerol (v/v), 100 mM NaCl, 1 M imidazole). Fractions (1 ml) were collected and the highest concentrations of protein were pooled.

Kinetic characteristics of each pyruvate kinase were determined using purified protein samples according to the method of Bergmeyer et al.,^38^ Assays to determine the enzyme kinetics for each PK under each condition were carried out in triplicate and analysed using GraphPad Prism using the Michaelis-Menten or Hill equation where appropriate.

### RNA extraction, semi-quantitative RT-PCR and qPCR analysis

Biomass of *S. coelicolor* came from liquid cultures and semi-quantitative RT-PCR was carried out according to the method of Clark and Hoskisson^7^. Total RNA was used as template for cDNA synthesis using a qPCRBIO cDNA synthesis Kit (PCR Biosystems). All cDNA samples were diluted to a concentration of 10 ng/μl with each qPCR reaction containing 10 ng of cDNA (1 μl) and were then mixed with 10 μl MasterMix (Kit 2x qPCRBIO SyGreen Mix Lo-ROX from PCRBIOSYSTEMS), 2.5 μl of each primer to a final reaction volume of 20 μl using the Corbett Research 6000 (Qiagen).

## Acknowledgements

We would like to thank the Scottish Universities Life Science Alliance (SULSA) for BioSkape PhD funding to JKS, The Mac Robertson Travelling Scholarship awarded to JKS, and the Natural Environment Research Council for Grant NE/M001415/1 to PAH. We would also like to thank Professor Ian Henderson and Dr Faye Morris (University of Birmingham) for the gift of the *E. coli* Keio collection mutants and Professor David Hodgson for helpful discussions.

